# Stimulation of zona incerta selectively modulates pain in humans

**DOI:** 10.1101/2020.09.02.277921

**Authors:** Charles W Lu, Daniel E Harper, Asra Askari, Matthew S Willsey, Philip P Vu, Andrew D Schrepf, Steven E Harte, Parag G Patil

## Abstract

Stimulation of zona incerta in rodent models has been shown to modulate behavioral reactions to noxious stimuli. Sensory changes observed in Parkinsonian patients with subthalamic deep brain stimulation suggest that this effect is translatable to humans. Here, we utilized the serendipitous placement of subthalamic deep brain stimulation leads to directly investigate the effects of zona incerta stimulation on human pain perception. We found that stimulation at 20 Hz, the physiological firing frequency of zona incerta, reduces experimental heat pain by a modest but significant amount, achieving a 30% reduction in one fifth of implants. Stimulation at higher frequencies did not modulate heat pain. Modulation was selective for heat pain and was not observed for warmth perception or pressure pain. These findings provide a mechanistic explanation of sensory changes seen in subthalamic deep brain stimulation patients and identify zona incerta as a potential target for neuromodulation of pain.

## Introduction

Deep brain stimulation (DBS) has been used for treatment of pain since the 1970s. Stimulation of classical targets—sensory thalamus, periaqueductal gray, and periventricular gray matter—is often able to provide pain relief, although longterm success of these interventions varies widely across etiologies. DBS treatment of phantom limb pain, failed back surgery syndrome, and trigeminal neuropathy is frequently successful; however, outcomes for other etiologies, including stroke, peripheral neuropathy, and brachial plexus injury, tend to be less satisfactory ^1-3^. The many pain patients for whom conventional DBS remains ineffective highlight the need for new targets of neuromodulation.

Emerging evidence points to zona incerta, a heterogeneous region of cell bodies and fibers dorsal to subthalamic nucleus, as a promising new target for pain neuromodulation. The region receives direct spinothalamic input and projects GABAergic efferents to ventromedial thalamus, which integrates cortical and spinothalamic inputs (Figure 1) ^4,5^. In a rodent model of central pain, zona incerta was shown to act as a feedforward inhibitor of pain perception ^6^. The same group later demonstrated that 50-60 Hz stimulation of zona incerta reduces hyperalgesia in a rat model of neuropathic pain, providing a proof of concept for analgesic zona incerta DBS ^7^. More recent work has established compelling causative links between GABAergic output from zona incerta, neuropathic pain, and neuromodulatory relief of hyperalgesia and allodynia ^8-10^.

**FIGURE 1.**
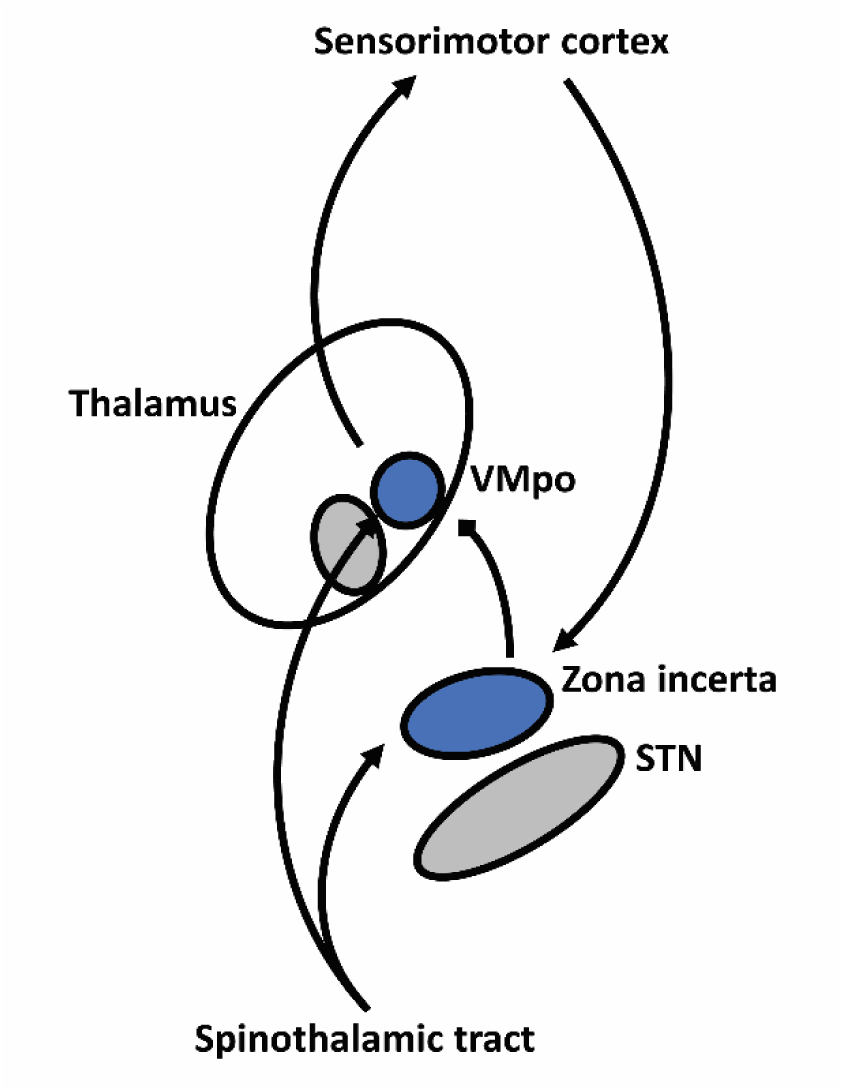
Feedforward inhibition of thalamic pain processing by zona incerta. VMpo: ventromedial posterior nucleus.

Experimental findings in rodent models have been corroborated by observations of sensory changes in human patients receiving subthalamic DBS. Although subthalamic DBS for Parkinson disease nominally targets the subthalamic nucleus, a large body of work has shown that most active contacts are located at or above the dorsal border of the subthalamic nucleus—a region directly adjacent to and overlapping zona incerta ^11,12^. While the intervention is best recognized for its suppression of parkinsonian motor symptoms, it is also known to have substantial therapeutic effects on pain and sensation ^13-17^. Multiple studies show that this effect is not explained by motor improvements alone ^18,19^, indicating an independent mechanism by which subthalamic stimulation ameliorates pain symptoms. Taken together, these observations strongly suggest that stimulation of zona incerta modulates pain perception.

In this study, we utilized the serendipitous placement of subthalamic DBS leads in patients with Parkinson disease to directly evaluate the effects of zona incerta DBS on human perception of experimental heat and mechanical pain. A broad set of stimulation parameters were tested and then prospectively validated in independent cohorts, showing that stimulation at physiological frequency modulated perceived heat pain by a modest but significant amount. Modulation was specific to heat pain and did not significantly alter perception of non-painful heat or mechanical pain. These findings provide a mechanistic explanation of sensory changes seen in subthalamic DBS patients and identify zona incerta as a potential target for neuromodulation of pain.

## Results

### Subjects

Subjects scored an average of 3.1 points (standard deviation of 2.5 points) on the Geriatric Depression Scale Short Form, with one subject (at 11 points) exceeding the 10-point cutoff for depression risk (included in analysis) (Table 1). One subject was a non-responder to hot stimuli up to 45 °C and was excluded from all analyses. Results were collected unilaterally on one subject due to scarring on one arm from previous traumatic injury. Unilateral data from one subject was excluded due to misinterpretation of subject instructions. No subjects reported pre-existing pain in the areas examined in this study.

**Table 1.** Subject characteristics.

| Age | Sex | GDSS | Duration of DBS,<br>months | Experiment | Data<br>inclusion |
| --- | --- | --- | --- | --- | --- |
| 62 | M | 4 | 55 | Exploratory | Bilateral |
| 58 | M | 2 | 18 | Exploratory | Bilateral |
| 62 | M | 2 | 64 | Exploratory | Excluded |
| 67 | M | 2 | 13 | Exploratory | Bilateral |
| 58 | F | 1 | 65 | Exploratory | Bilateral |
| 71 | M | 2 | 59 | Exploratory | Bilateral |
| 73 | M | 2 | 7 | Exploratory | Unilateral |
| 45 | M | 1 | 22 | Validation | Bilateral |
| 62 | F | 11 | 6 | Validation | Bilateral |
| 60 | M | 3 | 40 | Validation | Bilateral |
| 73 | F | 2 | 42 | Validation | Bilateral |
| 66 | M | 2 | 38 | Validation | Unilateral |
| GDSS: Geriatric Depression Scale Short Form score. Duration of DBS measured from time of implant to time of study. |  |  |  |  |  |

### Zona incerta DBS modulates heat pain

Zona incerta DBS with conventional 130 Hz stimulation decreased perceived heat pain by 0.71 points (*p*=0.01, n=99) on the visual analog scale (Figure 2). Low frequency 20 Hz stimulation reflecting physiological firing of zona incerta also reduced pain elicited by hot stimuli (−0.78 points; *p*=0.005). DBS of any frequency did not appear to significantly affect perceived pain from warm stimuli (20 Hz, *p*=0.23; 60 Hz, *p*=0.63; 130 Hz, *p*=0.35; n=99 for all comparisons) or mechanical pain thresholds (20 Hz, *p*=0.70; 60 Hz, *p*=0.15; 130 Hz, *p*=0.19; n=99 for all comparisons).

**FIGURE 2.**
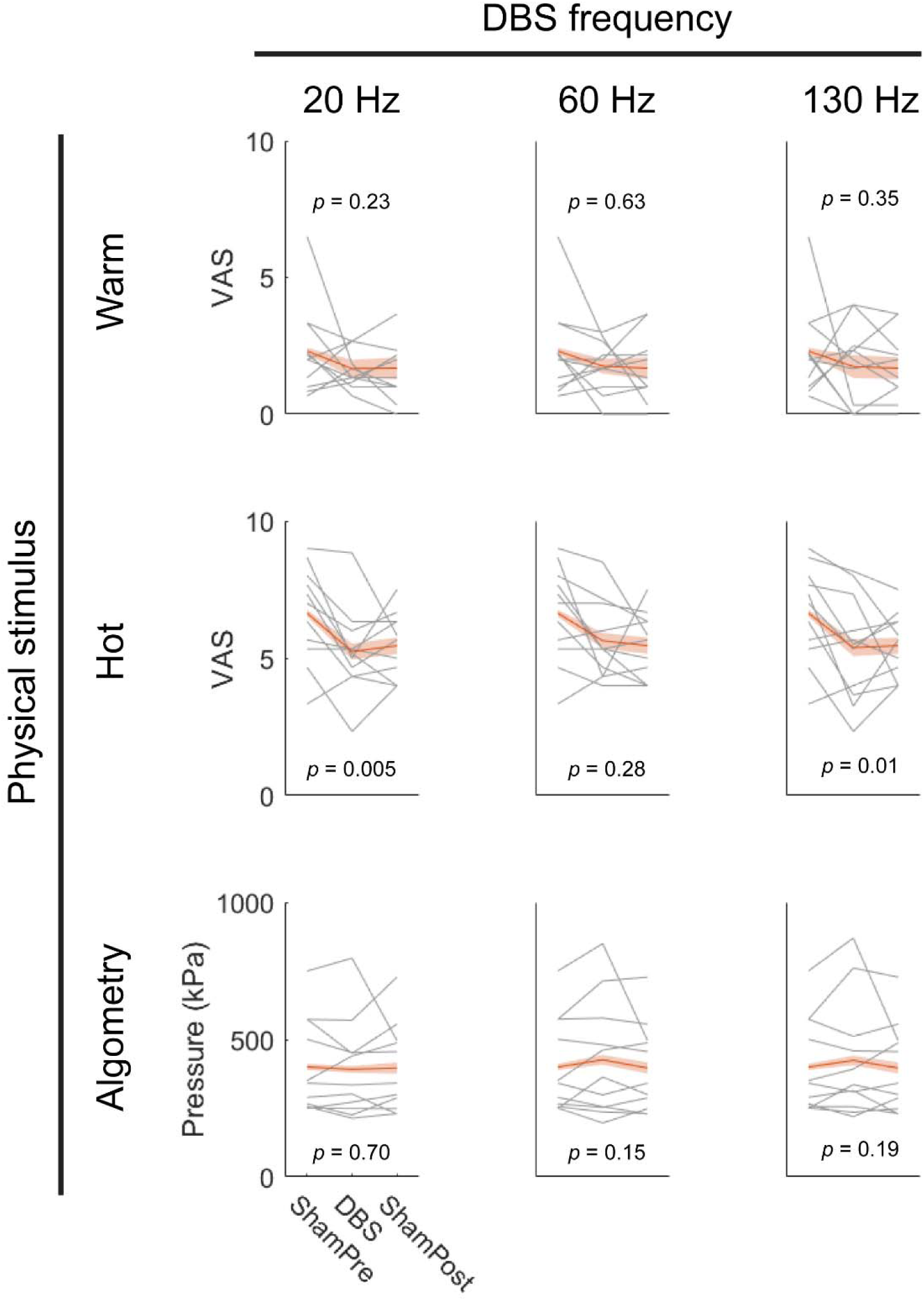
Effects of DBS on perceived pain from warm stimuli, hot stimuli, and mechanical pressure. Gray lines show mean pain scores across arm sites for each subject-implant. Red lines show average across implants with standard error of the mean shaded. *n* = 99 trials for all analyses shown, with sham trials shared across DBS frequencies. VAS: visual analog scale.

### Validation of 20 and 130 Hz stimulation

Due to the small effect sizes observed and multiple comparisons made in the exploratory experiments, the effects of 20 and 130 Hz stimulation on heat pain were measured in an independent set of nine implants (five subjects) to confirm results. This group also received an additional sham trial. In this cohort, 20 Hz stimulation reduced heat pain by 0.51 points (*p*=0.006, n=96), confirming the original observation of this effect (Figure 3a). Stimulation at 130 Hz also reduced heat pain by 0.27 points but did not reach significance (*p*=0.16, n=96). As the validation experiments did not incorporate wash-in time, results also indicate that neuromodulation of heat pain by zona incerta DBS takes rapid effect.

**FIGURE 3.**
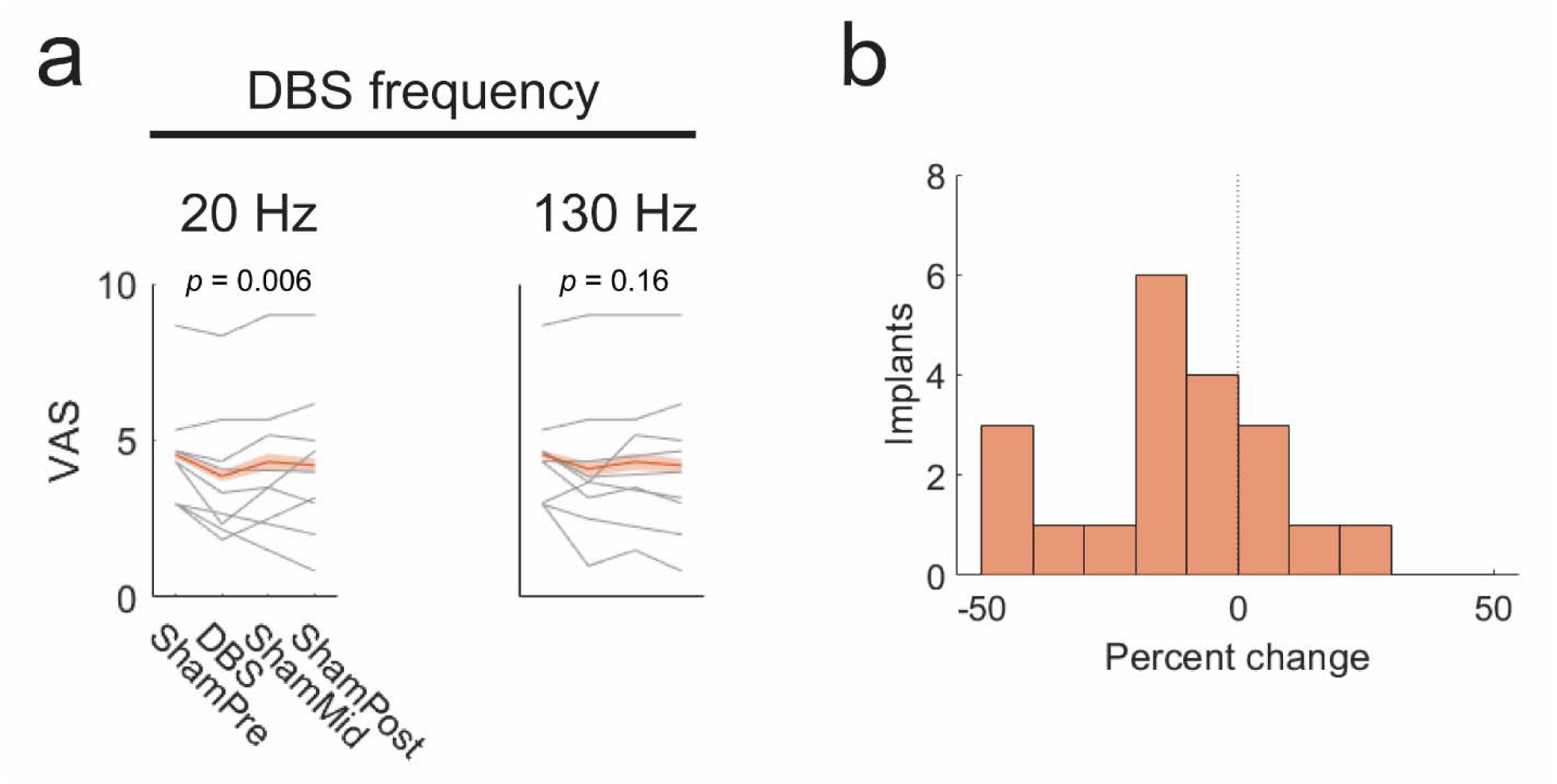
Effects of 20 and 130 Hz DBS on perception of heat pain. a. Effects of DBS on perceived pain from hot stimuli. Gray lines show mean pain scores across arm sites for each subject-implant. Red lines show average across implants with standard error of the mean shaded. *n* = 96 trials for both analyses, with sham trials shared across DBS frequencies. b. Distribution of percent change in heat pain with 20 Hz DBS. *n* = 20 implants.

To quantify the effects of 20 Hz stimulation on heat pain, data from exploratory and validation experiments were combined. Analysis of each subject-implant’s percent improvement from averaged sham score revealed that stimulation achieved pain reduction of 30% or more in 20% of implants. The mean and median effects of stimulation on heat pain were-11.8% and-11.3%, respectively, with a standard error of 4.5% (Figure 3b).

## Discussion

This study evaluates a new form of DBS for pain and demonstrates that stimulation of zona incerta achieves a modest but significant analgesic effect in human subjects. We further demonstrate that analgesia is best achieved using stimulation at low frequency. This phenomenon was selective to perception of heat pain and did not affect perceived intensity of warm or mechanical stimuli. The results of this study are the first to confirm that stimulation of zona incerta modulates evoked pain perception in humans. This follows a compelling body of work in rodent models, which have demonstrated both behavioral manifestations and mechanistic explanations of pain modulation by excitation of zona incerta ^7,8^. Human translation of zona incerta DBS is an important step to better qualify the perceptual effects of zona incerta neuromodulation and lays a foundation for further optimization of analgesic DBS.

Observation that analgesia is best achieved with stimulation at physiological frequency is a notable finding. While stimulation at conventional DBS frequencies has been hypothesized to act as an informational lesion ^20^, stimulation at physiological firing rates may act to increase activity in zona incerta, which has been shown by rodent studies to impart analgesic effect ^8^. A parsimonious interpretation of the findings is that analgesic stimulation acts by increasing GABAergic output from zona incerta to sensory thalamus. Importantly, we also show that the effects of stimulation appear specific to heat pain; perception of non-painful warm stimulation and mechanical pain thresholds were not altered by DBS. However, zona incerta is known to project widely across the brain ^5^, and potential relevant off-target effects were not investigated in this study, nor were other pain modalities.

There are differences in findings between this study and previous rodent studies. Most notably, this study did not identify any significant effects of zona incerta DBS on mechanical pain thresholds, while hind paw withdrawal thresholds were seen to increase in rodent models of neuropathic pain ^7,8,10^. Although unexpected, this may arise from a variety of differences between our study and those performed in rodent models. Primarily, pain in our patients did not arise from clinically relevant sources of neuropathic pain. Additionally, Parkinson disease is known to cause a broad but inconsistent and poorly understood constellation of sensory abnormalities ^21^, introducing an important confounder. Performing this study in humans, however, allowed for the first experiment to directly assess the effects of zona incerta stimulation on perceived pain intensities, rather than noxious withdrawal thresholds. Other human studies describing the sensory effects of nearby subthalamic stimulation differ on whether mechanical pain thresholds are modified by stimulation ^17,18^. However, the mechanistic pathway of these effects may also be distinct from that of DBS at zona incerta ^22^.

Critically, interpretation of these results must acknowledge that targeting of zona incerta in this study is inherently imprecise. While DBS leads for Parkinson disease are placed to activate dorsolateral subthalamic nucleus, the portions of zona incerta connected to the spinothalamic tract and sensory thalamus are found in ventral zona incerta, which is located medial to the dorsolateral horn of subthalamic nucleus ^5^. As such, optimal activation of the target region could not be guaranteed. The imprecise nature of stimulation targeting may account for some of the large variations in effect size observed across subjects shown in Figures 2 and 3.

Despite this limitation in study design, stimulation of zona incerta elicited a statistically significant and reproducible effect on perceived heat pain, identifying zona incerta as a strong candidate for neuromodulation of pain. Although clinical translation of this intervention requires substantial additional work, these findings provide a compelling explanation of how subthalamic DBS modulates pain-related symptoms observed in Parkinson patients and present clear avenues for optimization. Foremost, explicit targeting of ventral zona incerta, medial to the subthalamic DBS targets employed here, has potential to markedly improve both consistency and magnitude of the analgesic effect. Our finding that stimulation at physiological frequencies is effective also motivates further investigation of other low frequency stimulation paradigms and physiologically inspired patterns. More immediately, these results can be used to inform programming for the large population of subthalamic DBS patients presenting with pain. As we advance our understanding of zona incerta, further research in this direction is warranted, particularly to examine effects on clinically relevant etiologies of pain and sustainability of effects over longer time periods.

## Methods

### Subjects

Outpatient experiments were performed with 9 male and 3 female patients previously implanted with subthalamic DBS leads for treatment of Parkinson disease at the study institution. Patient selection criteria for subthalamic DBS at the institution have been described previously ^23,24^. All patients were implanted with Medtronic (Dublin, Ireland) DBS leads, model 3389, with guidance by 3T magnetic resonance imaging (MRI), stereotactic navigation, and microelectrode recording. Subjects were implanted at least six months prior to the study and had stable, effective programming parameters. Prior to study, outpatient subjects were screened for depression using the Geriatric Depression Scale Short Form ^25^. The study was approved by the Institutional Review Boards of the University of Michigan Medical School, experiments were performed in accordance with institutional guidelines and national regulations, and all participants provided individual informed consent.

### DBS lead placement

DBS targets were initially assigned from indirect targeting (12 mm lateral, 3 mm posterior, and 4 mm inferior to the mid-commissural point) and adjusted with direct visualization of the ventral border of subthalamic nucleus on 3T MRI (Philips Achieva 3T; Philips, Amsterdam, Netherlands). Microelectrode signals were recorded with a Neuroprobe amplified by a Neuro Omega system (Alpha Omega, Alpharetta, GA). Recordings were performed from 15 mm above to 5 mm below the planned target. An experienced electrophysiologist identified the location subthalamic nucleus during surgery. DBS leads were inserted with the tip near the electrophysiologically defined ventral border of subthalamic nucleus. Pulse generators were implanted and connected to DBS leads within 14 days of lead implantation. High-resolution computed tomography scans (GE HD750; General Electric, Boston, MA) were acquired two to four weeks after surgery to verify lead locations.

### Estimation of stimulation sites

Post-operative computed tomography scans were co-registered with magnetic resonance images using commercial software (Analyze; AnalyzeDirect, Overland Park, KS). Coordinates of DBS contacts were recorded, alongside coordinates of the subthalamic nucleus midpoint and its mediolateral, dorsoventral, and anteroposterior spans. Coordinates were linearly transformed into a common space with a shared orientation (left brain) and subthalamic nucleus midpoint. Coordinates for each contact were then scaled according to the size of its corresponding subthalamic nucleus to preserve relative anatomical locations. Scaled leads were then visualized within a representative magnetic resonance image to approximate the anatomical location of contacts used to deliver stimulation.

### Deep brain stimulation

Neuromodulation of zona incerta was achieved using the implanted DBS leads and pulse generators by delivering stimulation to the DBS contact closest to 1.5 mm above the dorsal border of electrophysiological subthalamic nucleus. Three different stimulation frequencies were used: 20 Hz, 60 Hz, and 130 Hz; which reflect the frequency of observed human ZI activity ^26^, the frequency of analgesic ZI stimulation in rats ^7^, and the frequency of conventional subthalamic DBS stimulation, respectively. Stimulation was delivered contralateral to the side of sensory testing with 60 µs charge-balanced pulses and voltage at 0.5 V below sensory threshold at 130 Hz, with a maximum of 2.0 V. Stimulation settings were set by an experienced clinician using a clinical programmer, with subject and experimenter blinded to stimulation settings. Estimated simulation sites for subjects in the Exploratory experiments (see *Experiment design*) are shown in Figure 4a.

**FIGURE 4.**
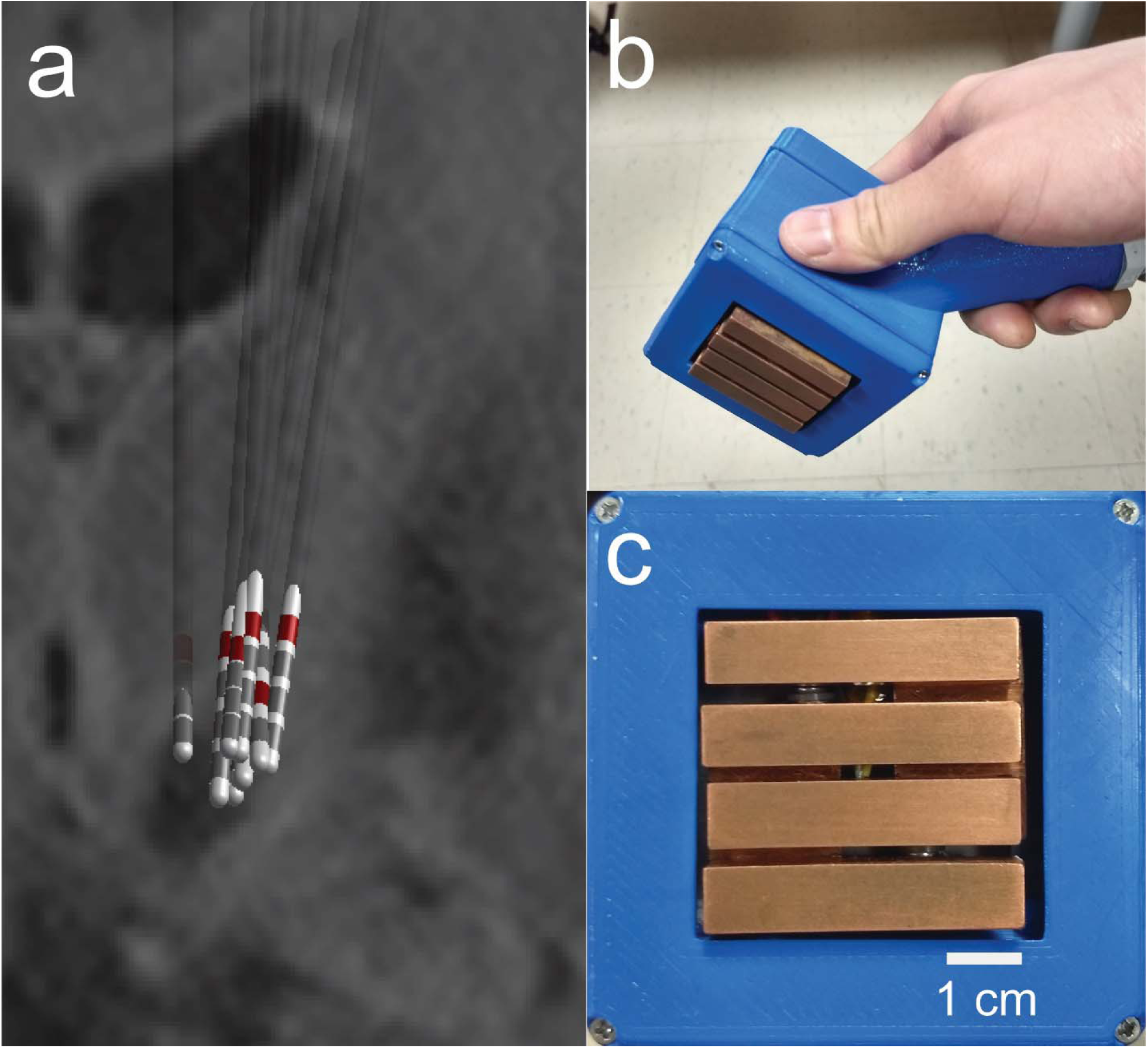
Deep brain stimulation sites and thermal stimulation device. a. Estimated lead locations in subjects participating in exploratory experiments juxtaposed against a representative magnetic resonance image. Active contacts used in the study are indicated in red. b. Device used to produce thermal stimuli. c. Detail of device contact surface.

### Thermal stimulation

A custom device was used to provide thermal stimuli. The contact surface is composed of four parallel copper bars (9×10 mm surface), shown in Figure 4b and 4c, with temperature controlled by Peltier devices. Two distinct stimuli were produced with the device: nonpainful *Warm* stimuli were produced by setting bars to 39 °C; painful *Hot* stimuli were achieved by setting bars to 45 °C, or the highest temperature tolerable by the subject (always greater than 41 °C).

Thermal stimuli were applied to three sites along the volar forearm: proximal aspect, midpoint, and distal aspect, centered along midline. Each thermal stimulus was tested once at each site for each DBS setting. Application of thermal stimuli followed the sequence listed above with at least 30 seconds of rest between successive applications. After application of each thermal stimulus, patients were asked to separately rate the intensity (See Supplementary Fig. S1 online) and pain of each thermal stimulus on a 10-point scale, with 0 signifying “no sensation/pain” and 10 signifying the “most intense/painful sensation imaginable.”

### Mechanical stimulation

Algometry was performed using an Algometer type II device (SBMEDIC Electronics, Solna, Sweden) upon the belly of the extensor digitorum muscle. Three measurements of pressure pain threshold were performed for each test case at locations roughly 1 cm apart. Measurements across test cases were performed at overlapping but non-identical locations.

### Experiment design

#### Exploratory experiments

A set of experiments evaluating the effects of 20, 60, and 130 Hz zona incerta stimulation on perceptions of warm, hot, and mechanical stimulation was performed bilaterally with seven outpatient subjects (Figure 5a).

**FIGURE 5.**
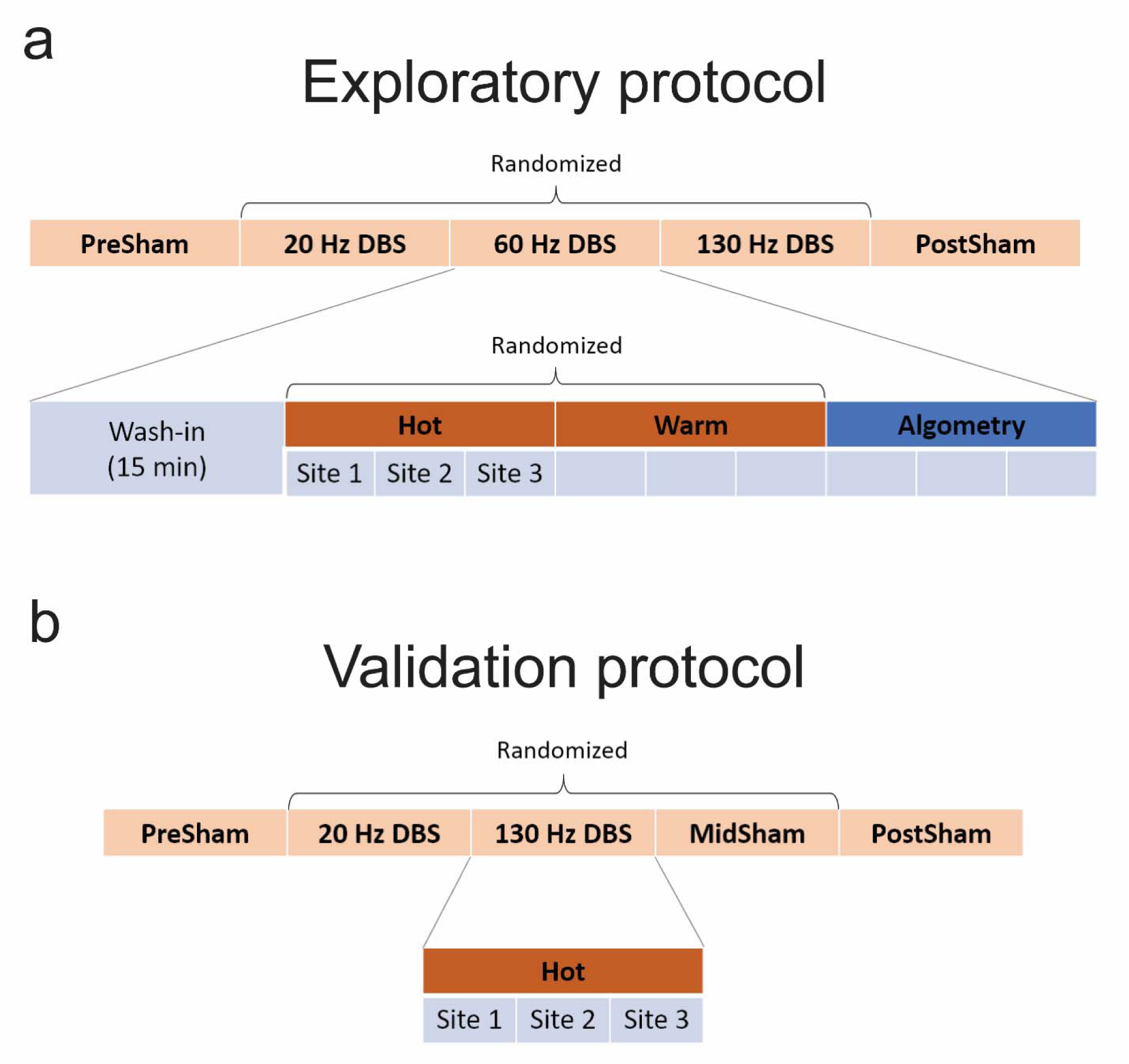
Experimental protocols. a. Protocol for initial exploratory trials. b. Protocol for validation trials including additional sham trial. Order of randomized DBS settings were hidden from both subject and experimenter. Order of randomized thermal stimuli were hidden from subject.

Each DBS setting was applied for 15 minutes. The first ten minutes involved no sensory testing to allow for wash-in of potential slow-acting effects. The last five minutes were used to perform sensory testing. Thermal stimuli (warm, hot) were tested in randomized order, counterbalanced across DBS settings. Algometry was performed after thermal testing. Patients were tested unilaterally on one side first, then the other. The first and last DBS settings were sham stimulation. The order of 20, 60, and 130 Hz stimulation was randomized. The experimenter was blinded to stimulation frequency. Test subjects were blinded to stimulation setting (sham and frequency) and thermal stimulation paradigm.

#### Validation experiments

A second set of experiments was performed on an independent cohort of five subjects to evaluate the effects of 20 and 130 Hz zona incerta stimulation on perceived heat pain (Figure 5b).

Validation experiments followed a similar protocol to that of exploratory experiments, described above. However, no wash-in time was provided. Instead, thermal stimulation followed immediately (within five minutes) after application of DBS settings. Only hot stimuli were used for sensory testing. In addition, 60 Hz stimulation was replaced with another period of sham stimulation for three of the five subjects. Subjects remained blinded to all deep brain stimulation and sensory stimuli settings. Experimenter was blinded to order of stimulation parameters (including the additional sham period).

#### Statistical analysis

A mixed linear model controlling for differences in patient baselines (*β*_*subject*_) and habituation over time (*β*_*habit*_) was used to determine the effect and significance of each intervention by zona incerta stimulation (*β*_*DBS*_).

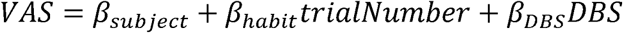

Each implant was treated as an individual subject during statistical analysis. As such, results obtained from contralateral sides of bilaterally tested subjects were assumed to be independent and have different baselines. Sham stimulation trials were shared across interventions. Results of interest were validated with an independent set of subjects, in lieu of adjusting for multiple comparisons.

## Acknowledgements

The authors would like to thank Adam Davis for his assistance in designing and constructing the thermal stimulation device. This study would not have been possible without the clinical experience and help of Kelly Lupo, Wilma Mackenzie, and Adam Matthews.

This work was supported by the A. Alfred Taubman Medical Institute, Ann Arbor, MI; the Coulter Foundation, Ann Arbor, MI; the STIM (Surgical Therapies Improving Movement) Program, Ann Arbor, MI; and the University of Michigan Medical School, Ann Arbor, MI.

The authors have no conflicts of interest to disclose.

## Author Contributions

CWL: Conception, Design, Acquisition, Analysis, Interpretation, Software, Drafting; DEH: Conception, Design, Interpretation; AA: Acquisition, Analysis; MSW: Conception, Acquisition; PPV: Acquisition; ADS: Design, Interpretation; SEH: Conception, Design, Interpretation; PGP: Conception, Design, Interpretation.

## Additional Information

The authors declare no competing interests. Data from this study are available upon reasonable request to the corresponding author.

## Supplement

**SUPP FIGURE 1:**
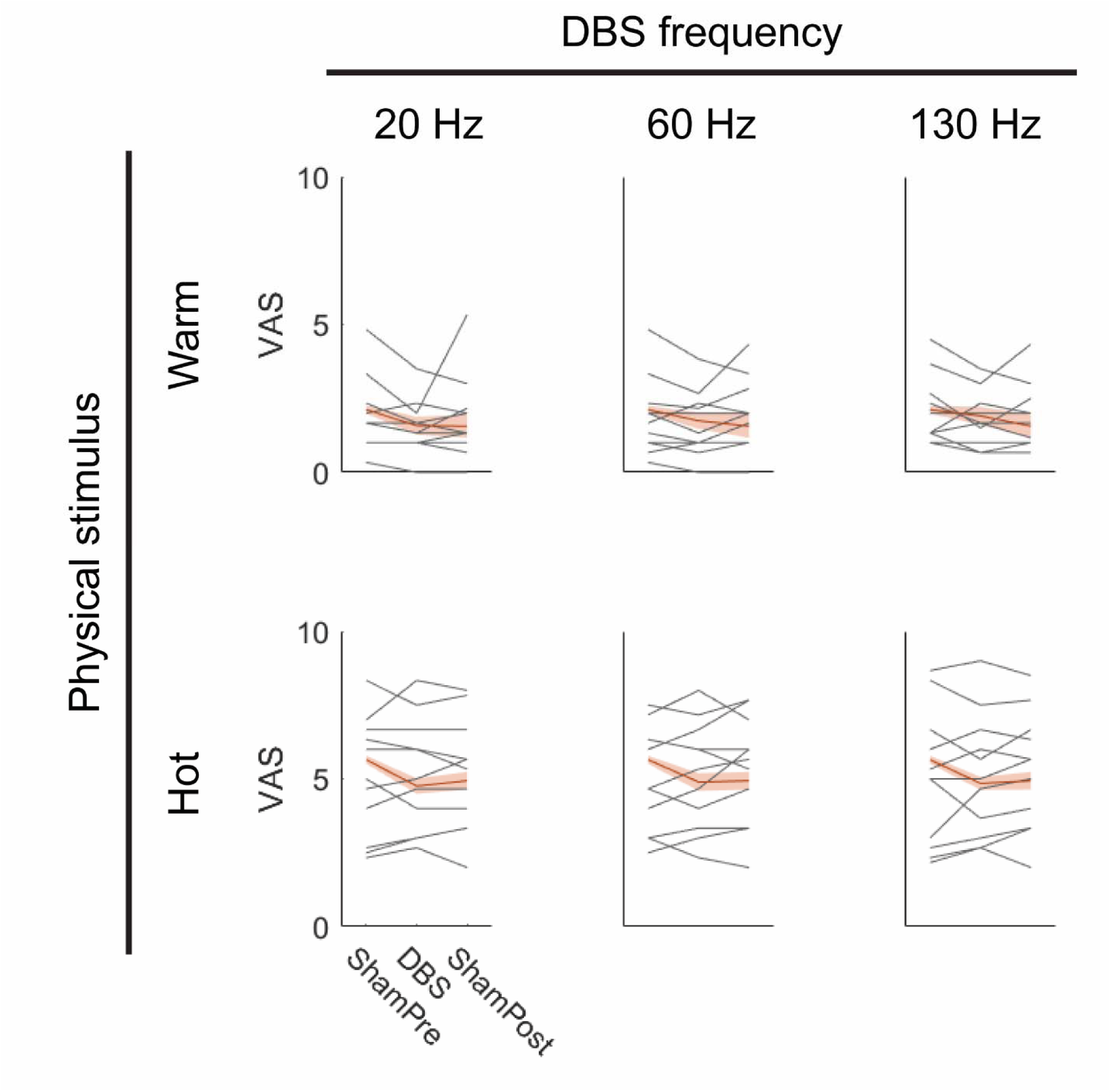
Effects of DBS on perceived intensity from warm and hot stimuli. Gray lines show mean intensity scores across arm sites for each subject-implant. Red lines show average across implants with standard error of the mean shaded. *n* = 99 trials for all plots, with sham trials shared across DBS frequencies.

